# Resting brain activity emerges from wave propagating along spatiotemporal varying hyper-structural connectome

**DOI:** 10.1101/2021.10.11.464009

**Authors:** Yanjiang Wang, Jichao Ma, Qingwei Meng, Xue Chen, Chunyu Du

**Author notes:** Corresponding author. (Y. Wang).

## Abstract

How spontaneous brain activities emerge from the structural connectivity (SC) has puzzled researchers for a long time. The underlying mechanism still remains largely unknown. Previous studies on modeling the resting-state human brain functional connectivity (FC) are normally based on the relatively static structural connectome directly and very few of them concern about the dynamic spatiotemporal variability of FC. Here we establish an explicit wave equation to describe the spontaneous cortical neural activities based on the high-order hypergraph representation of SC. Theoretical solution shows that the dynamic couplings between brain regions fluctuates in the form of an exponential wave regulated by the spatiotemporal varying Laplacian of the hyper-structural connectome (*h*SC), which orchestrates the cortical activities propagating in both space and time. Ultimately, we present a possible mechanism of how negative correlations emerge during the fluctuation of the hypergraph Laplacian of SC, which helps to further understand the fundamental role of SC in shaping the entire pattern of FC with a new perspective. Comprehensive tests on four connectome datasets with different resolutions confirm our theory and findings.

## 1. Introduction

The human brain is highly active all the time, being at rest or task-evoked, with signals propagating between sets of brain areas along structural pathways (e.g., the cortical white matters). Diffusion magnetic resonance imaging (dMRI) techniques show that the brain regions are sparsely anatomically connected through white matter fibers, while studies on functional MRI (fMRI) estimated from the blood oxygenation level-dependent (BOLD) signals reveal that high correlations can also be observed between brain areas without directly anatomical linkage. Although mounting evidence supports the view that FC is determined by the underlying SC [1–3], the mechanism behind the shaping of the entire pattern of FC especially the emergence of negative correlations is still poorly understood. To this end, a vast body of research has been conducted to disclose this riddle by modeling FC using SC since the last decade. For example, using complex network and graph modeling methods [4, 5], some studies indicate that a wide variety of structural network measures, such as shortest path length and steps [6, 7], search information and path transitivity [8, 9], as well as node degree product [10, 11], etc., are all related to the presence of FC, but to a limited extent. Whereas computational neural mass models (NMMs) [12–15] aim to uncover the inherent complex neural dynamics between neuronal populations in human brain, which entails a number of empirical parameters to be pre-specified or experimentally tuned, thus making the models highly intricate to handle. Meanwhile, some other methods that relate SC and FC using direct matrix mapping [7, 16, 17] or spectral graph mapping [18–20] are intensively explored as well, in which the FC matrix is simply formulated as the weighted sum of the SC matrix and its higher order indirect dependency matrices, with the weights being trained for each group of SC and FC matrices. These models, albeit show high accuracy with the trained dataset, will fail to generalize to other connectome datasets.

Recently, the graph Laplacian of the human connectome has been found to be able to capture the relationship between brain structure and function. The graph diffusion (GD) model [21, 22], which relies mainly on the graph Laplacian of the brain SC, has shown potential advantages in predicting FC with few model parameters. Nevertheless, the GD model lacks the ability to yield negative functional correlations and only a small part of FC between brain areas that anatomically unconnected can be predicted due to the sparseness of the graph Laplacian matrix of SC. To remedy the limitations of graph Laplacian, we extended the GD model in recent studies [23, 24] to two hypergraph based diffusion models, HGD and HpGD, based respectively on the hypergraph Laplacian and *p*-Laplacian [25–27] of human brain connectome and obtained much better results in modeling FC, which are even capable of modeling anti-correlations by embedding a matrix with each entry indicating the sign of the correlation between two brain areas.

In this study, we aim at unraveling how spontaneous brain activities from different areas are coupled and propagate in space and time by establishing a wave equation using the hypergraph Laplacian of SC. We first define the hypergraph representation of human brain SC concisely and then present the notation of hyper-structural connectome (*h*SC), i.e, the adjacency matrix of the hypergraph of SC, as well as the high-order representation of *h*SC (*h*^*m*^SC, with *m* being the number of order) derived from the incidence matrix of the hypergraph of SC to demonstrate how brain signals spread in a hyperpraph. We then elaborate the derivation of the wave equation to show how the brain FC are shaped including the emergence of negative correlations based on the time-varying hypergraph Laplacian of SC. Finally, we apply the wave equation to four extensively studied experimental connectome datasets with increasing resolutions to demonstrate its power in mapping human brain SC to FC during rest.

## 2. Hypergraph representation of SC

### 2.1 Hypergraph notation

Different from the definition of hypergraph for dynamic functional connectivity analysis in a previous study [28], in our approach, the hypergraph of SC is defined by ***H*_*g*_ = (*V*, *E*, *W*_*h*_)** with a set of *n* vertices representing the brain areas, *V* = {*v*_*i*_|*i*∈1,2,…,*n*}, and a set of *n* hyperedges with each containing all edges linking to the corresponding brain area, *E*={*e*_*j*_|*j*∈1,2,…,*n*}. The weight strength of each hyperedge is quantified by summing all the connecting edge weights contained in the hyperedge and denoted as *W*_*h*_ = {*w*_*hj*_|j∈1, 2,…*n*}. The relationship between *V* and *E* can be expressed as a |*V*| × |*E*| matrix ***H***, namely the incidence matrix of ***H***_***g***_, with entries *h*(*v*, *e*) = 1 if *v*∈*e* and 0 otherwise. The degree of a vertex *v*∈*V* is defined as 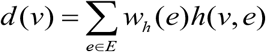, while the degree of a hyperedge *e* ∈ *E* is defined as *δ*(*e*)=Σ_*v*∈*V*_ *h*(*v*, *e*).

### 2.2 Hyper-structural connectome

From the above hypergraph definition, it can be inferred that a hyperedge of ***H***_***g***_ is capable of connecting more than two brain areas, and the incidence matrix of ***H***_***g***_, ***H***, can associate each hyperedge with brain areas belonging to the hyperedge. Therefore, a link between two brain areas that are not structurally connected but in the same hyperedge can be established via one intermediate area in the same hyperedge. As illustrated in Fig.1, a 9-node simulated brain graph network in which node V_1_ is initially connected to node V_2_ only, with its structural connectome rendered as a matrix shown below the graph (Fig.1 A). According to the definition of the hypergraph adopted in this study, 9 hyperedges can be defined initially in the hypergraph and, for visualizing clearly, only two hyperedges, e_1_ and e_2_, are labeled (Fig.1 B), where e_1_ contains nodes V_1_ and V_2_, e_2_ contains nodes V_1_, V_2_, and V_3_, i.e., in this case, V_1_, V_2_ and V_3_ belong to the same hyperedge. Then a link can be established between nodes V_1_ and V_3_ via V_2_, which in turn yields a new adjacency matrix of SC, termed hyper-structural connectome (*h*SC), as the matrix below the graph shown in Fig.1 C.

**Fig. 1.**
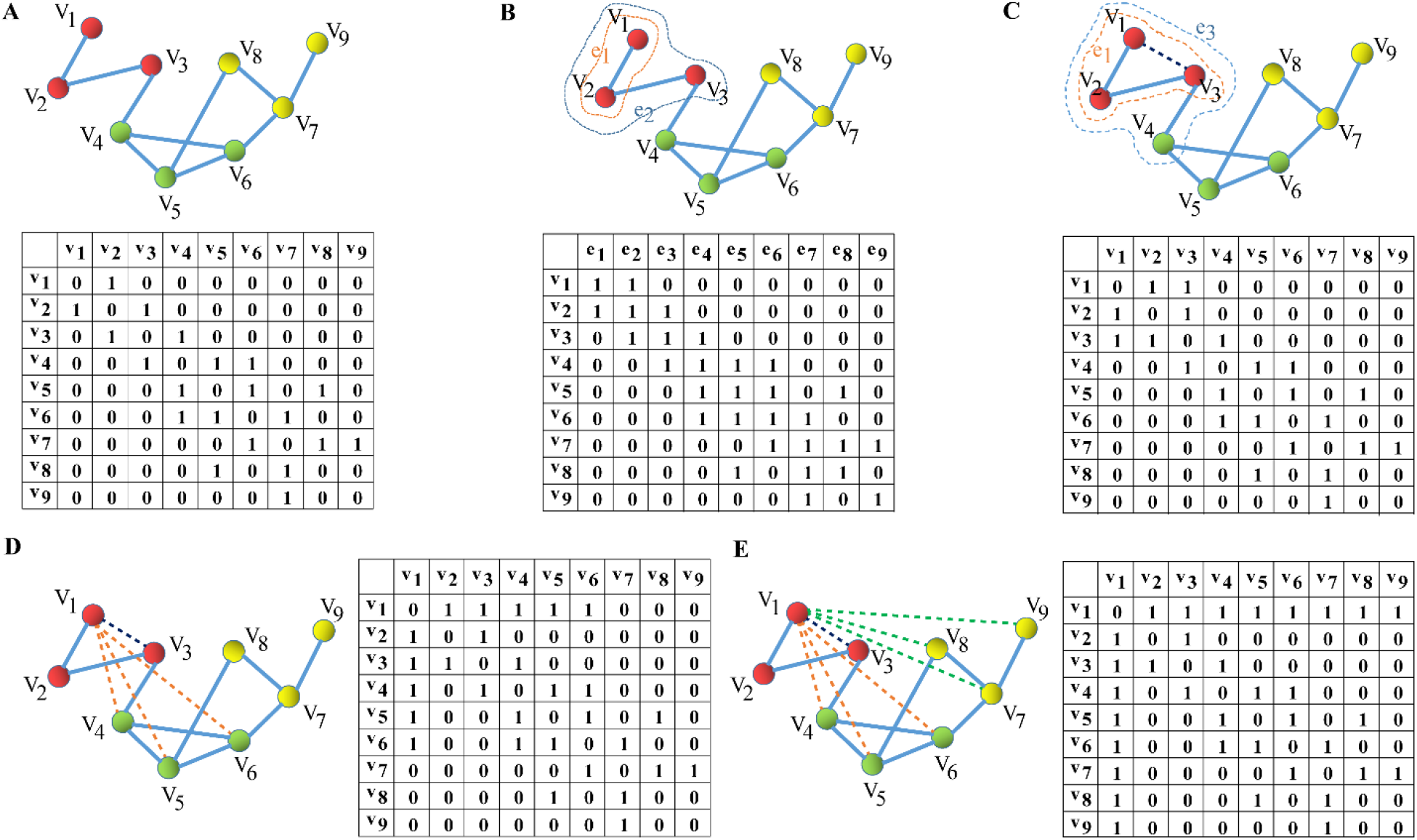
Illustration of the shaping of high-order of *h*SC from a simulated 9-node graph network. **(A)** A simulated 9-node brain graph and its adjacency matrix. **(B)** The hypergraph representation of the graph in (A) and its incidence matrix, in which, only two hyperedges, e_1_ and e_2_, are labeled. **(C)** The first-order of *h*SC of the hypergraph in (B), in which, node V_1_ is linked to V_3_. **(D)** The second-order of *h*SC (*h*^*2*^SC) yielded from the hypergraph in (C) together with its adjacency matrix, in which, node V_1_ is linked to V_4_, V_5_ and V_6_. **(E)** The third-order of *h*SC (*h*^*3*^SC) shaped from the hypergraph in (D) together with its adjacency matrix, in which, node V_1_ is linked to the remaining nodes V_7_, V_8_, and V_9_. Note that here we just take the signal spreading from node V_1_ to other nodes as an example.

Specifically, the hyper-structural connectome can be defined by a matrix mapping ***H***_***g***_: ***SC*→*hSC***, i.e., 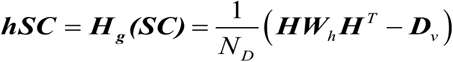, where ***D***_*v*_ denotes the diagonal matrix containing the vertex degrees, ***W***_***h***_ represents the diagonal matrix comprising the weights of hyperedges, while 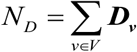 is a normalized factor that ensures the entries in *h*SC are less than one.

Similarly, the high order of *h*SC can be defined by a high-order matrix mapping, 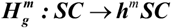, i.e., ***h***^***m***^***SC***=***H***_***g***_(…***H***_***g***_(***SC***))=***H***_***g***_^***m***^(***SC***), with *m* being the number of orders, *m* > 1. As will be clearly explained, the major distinction between *h*^*m*^SC and SC is that *h*^*m*^SC is in a state of flux both in space and time whereas SC is usually regarded as relatively fixed.

As shown in Fig. 1 C, the hypergraph will be redefined after V_1_ and V_3_ linked, in which hyperedge e_1_ contains nodes V_1_, V_2_, and V_3_, while e_3_ contains nodes V_1_, V_2_, V_3_, and V_4_. It should be noted that a link will be built between any two unconnected nodes connecting to a common node after the first-order mapping of *h*SC, for examples, node pairs between V_2_ and V_4_, V_3_ and V_5_, V_3_ and V_6_, etc., which are not displayed. Here we just take the signal spreading from node V_1_ to other nodes as an example. Next we can obtain the second-order of *h*SC from the updated hypergraph in Fig.1 C, in which V_1_ is linked to V_4_ via V_3_ (or V_2_), to V_5_ and V_6_ via V_3_ (Fig.1 D) and then the third-order of *h*SC (Fig.1 E), in which V_1_ will be linked to the remaining nodes V_7_, V_8_, and V_9_. If all nodes are taken together, the signals from each node will be coupled with each other and lead to the spread of signals across all the nodes very rapidly.

### 2.3 Hypergraph Laplacian of SC

Let ***D***_*e*_ denote the diagonal matrix containing the hyperedge degrees, similar to graph Laplacian, the normalized hypergraph Laplacian of SC can be defined as: 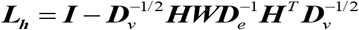 [25,26], where ***I*** indicates the identity matrix. Then the eigenvectors of ***L***_***h***_, representing the connectome harmonics of *h*SC, *ψ*_*j*_, *j*∈{*1*,…,*n*} can be obtained by solving the eigen-decomposition equation *L*_*h*_*ψ*_*j*_=*λ*_*j*_*ψ*_*j*_, *j*∈{*1*,…,*n*}, with *λ*_*j*_ being the corresponding eigenvalues of ***L***_***h***_, representing the frequencies of *h*SC [29–31].

## 3. Brain wave equation

Consider there are *n* brain areas and the dynamic cortical neural activities satisfy the following regular wave equation:

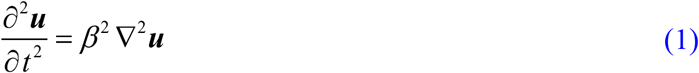

where ***u***=[*u*_1_, *u*_2_,…,*u*_*n*_]^*T*^, *u*_*i*_ = ***u***(*i*,*t*) refers to the neural activity signal of the *i*th brain area at time *t*, *T* indicates transpose, ∇^2^***u*** is the Laplace operator of ***u***, and *β* is the propagating factor.

Let *w*_*i,j*_ denote the connection strength between brain area *i* and *j* in *h*SC, with *w*_*i,j*_ satisfying: 0 ≤ *w*_*i,j*_ ≤ 1, *w*_*i,j*_ =*w*_*j,i*_, *w*_*i,i*_ = 0.

When the neurons in the *i*th brain area are firing, an influence of the *i*th area over the *j*th area occurs through signal propagating along the cortical pathways between the two areas, leading to a little fluctuation of the weight *w*_*i,j*_ varying with time *t*, denoted by Δ*w*_*i*,*j*_(*t*). Note that Δ*w*_*j,i*_(*t*) can be either positive or negative, representing the increase or decrease of the connection strength, respectively. Usually, the magnitude of Δ*w*_*j,i*_(*t*) varies linearly within a short period of time *t* and can be modeled as

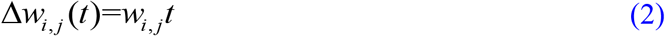

Similarly, Δ*w*_*j,k*_(*t*) can be expressed as

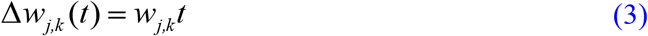

Then, the total first-order influence of area *i* over all the other areas amounts to

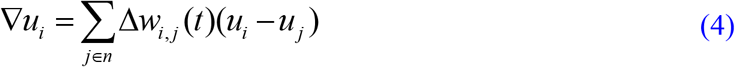

Likewise, the total first-order influence of area *j* over all the other areas is

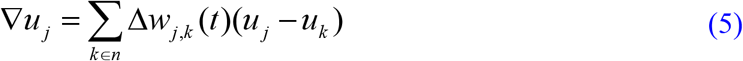

The above first-order process can be extended to the second order, i.e., the Laplace operator, of the *i*th area over the remaining areas, i.e.,

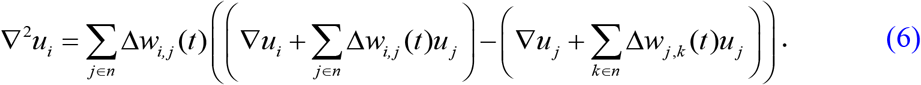

Substituting Eqs. (4) and (5) into Eq. (6), we obtain,

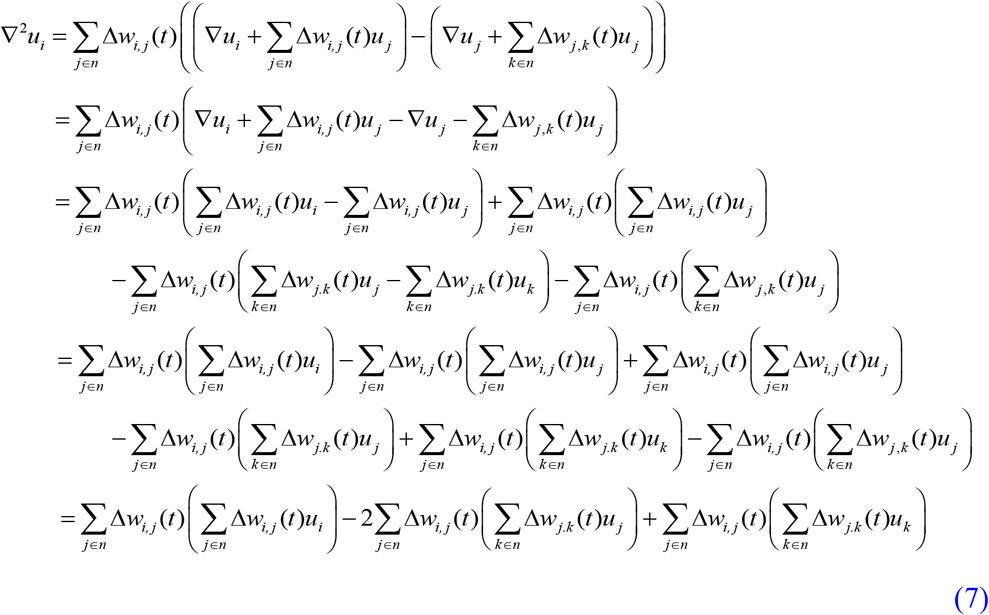

Substituting Eqs. (2) and (3) into Eq. (7), we obtain

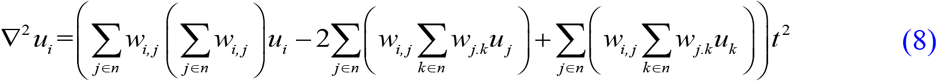

Considering all the *n* brain areas, we arrive at

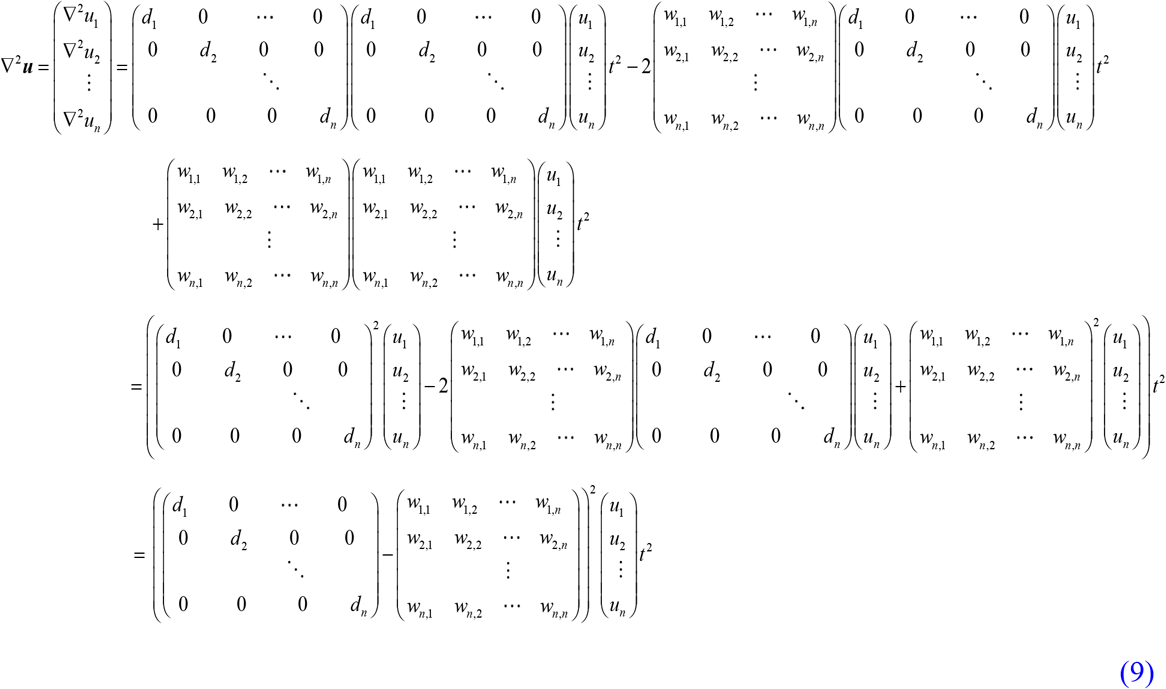

where 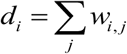 is the degree of area *i*. Define the degree matrix and adjacency matrix of *h*SC as ***D*** = *diag*(*d*_*i*_) and ***W*** = [*w*_*i,j*_], respectively, then we have

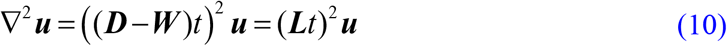

where ***L*** is the Laplacian of *h*SC, and the wave equation can be written as

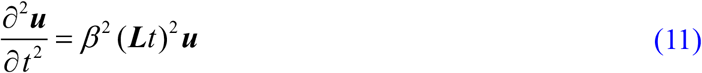

As mentioned above, since the connection strength between two brain areas might increase or decrease, causing the sign of Δ*w*_*i,j*_(*t*) unknown. Here, we make the assumption according to Hebbian learning rule [32], if the increase of the weight strength lasts for a long time, then long term potentiation (LTP) occurs and the two regions show positive correlation, otherwise long term depression (LDP) appears and the two regions yield negative correlation. Thus we modify the Laplacian matrix ***L*** as ***L***_***s***_ = ***L o S***, where ***S*** is a matrix with each entry indicating the sign of the correlation between two brain areas, ‘°**’** denotes the Hadamard product.

Then, the wave equation can be rewritten as

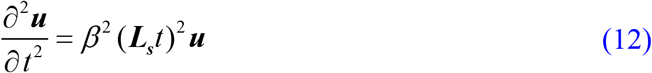

The solution of the above equation can be derived as (see *SM Note* 1)

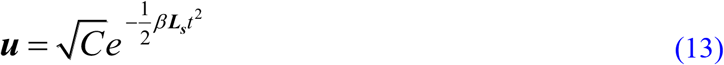

Factorizing matrix ***L***_***s***_ into its eigenvalues ***λ*** and eigenvectors ***U***, i.e., ***L***_***s***_=***UλU***^*T*^, the above equation becomes

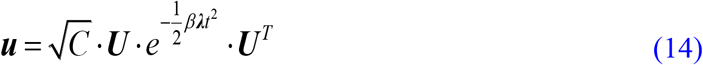

Then the dynamic couplings between brain areas can be quantified as the instantaneous autocovariance of ***u***, as follows,

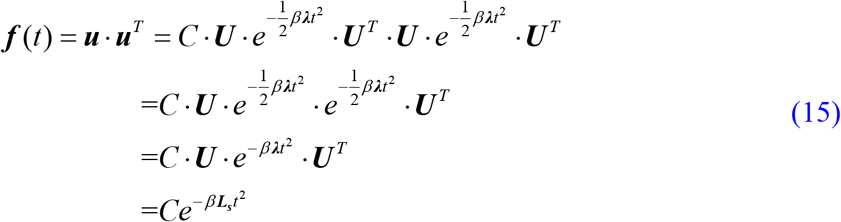

where *C* is a constant, keeping the strength of ***f***(*t*) within [0, 1].

## 4. Results

### 4.1 Connectome datasets

Four experimental datasets with 90, 246, 998, and 2514 ROIs (Regions of Interest) were exploited to evaluate the performance of the abovementioned wave equation and hypergraph model on SC-FC mapping, respectively, in which the SC matrices were derived from dMRI and tractography and the FC matrices were estimated using the resting-state fMRI data collected on a total of 170 subjects.

1. 90-ROI dataset The 90-ROI dataset was first reported in [21] and further studied in [7, 22–24], in which the structural and functional connectome data were obtained from 8 healthy adults and partitioned into 90 cerebral regions. The SC was obtained using probabilistic fiber tracts tracking and the connection weight between any two brain areas was estimated by the weighted sum of the fiber tracts going between them. The FC was measured by calculating the Pearson’s correlations between the BOLD time series recorded from each brain region.
2. 246-ROI dataset The original 246-ROI dataset is a multimodal MRI data that includes structural MRI (S-MRI), diffusion MRI (D-MRI) and resting-state functional MRI (R-fMRI), which were derived from 147 young healthy subjects at Beijing Normal University [33]. In our study, 145 subjects were chosen and the human Brainnetome atlases of 246 brain ROIs were employed (http://atlas.brainnetome.org) [34]. White matter tractography and probabilistic tracking method were performed to estimate the connection density between ROIs. The SC matrix of each subject was inferred through PANDA software (https://www.nitrc.org/projects/panda/), in which each entry represents the connection probabilities between ROIs. Those spurious weak connections whose probabilities are less than 1.0*e*-5 were removed. Besides, similar to the 90-ROI dataset, the resting-state FC matrices were also estimated using the Pearson’s correlations between the BOLD time series.
3. 998-ROI dataset The high resolution connectome dataset consisting of 998 ROIs has been extensively explored in many studies over the past decade [2, 4, 7, 8, 17], in which the structural and functional connectome data were obtained from 5 healthy right-handed male human participants adults and partitioned into 66 cerebral regions first and then subdivided into 998 ROIs. The SC matrices were defined by the number of fibers per unit surface connecting two regions (connection density). The FC matrices were also estimated using the Pearson’s correlations between the BOLD time series. For specific experimental details, see [4].
4. 2514-ROI dataset The very high resolution dataset with 2514 ROIs can be downloaded from the Brain Hierarchical Atlas NITRC page (http://www.nitrc.org/frs/?group_id=964), in which the SC matrix and resting-state FC matrix were derived from 12 healthy human subjects using diffusion tensor imaging (DTI) and tractography based on a parcellation of 2514 gray matter cortical areas. The SC matrices were obtained by quantifying the number of fiber tracts connecting each pair of ROIs. The FC matrices were also estimated using the Pearson’s correlations between the BOLD time series. For more experimental details, see [35].

The matrices’ elements of all the datasets except the 2514 ROIs are arranged such that the right hemisphere is in the upper left quadrant, left hemisphere in the lower right quadrant, and interhemispheric connections in off-diagonal quadrants. For clear visualization, the FC matrix of 2514 ROIs is modularized with hierarchical agglomerative clustering (HRC) and the SC matrix is binarized in the same order as those in the rsFC [35]. The mean measured FC and SC matrices of the four datasets are shown in Fig. S1. Note that we didn’t perform any further processing to the connectomes data during mapping SC to FC, with all the negative connections reserved in FC and different connection measures adopted in SC.

### 4.2 Parameters setting

Unlike previous eigen-decomposition based approaches [20], in which a number of parameters need to be learned from the measured FC, no parameters need training in our approach. There are only two free factors in Eq.(15), the constant *C*, which keeps the magnitude of the ***f***(*t*) within bounds, can be initially set to be 1, while the propagating factor *β*, controlling the time width of ***f***(*t*), can be chosen according to the number of brain areas parcellated. These two parameters have no much influence on the Pearson correlations between the measured FC and ***f***(*t*), allowing for determining the resting-brain FC from ***f***(*t*) in terms of the Pearson correlation accurately.

### 4.3 Structure-function mapping using Laplacian of *h*^*m*^SC

Firstly, for each subject of the four datasets, we calculate the simulated instantaneous functional correlation ***f***(*t*) between brain areas using Eq. (15), and the performances are assessed with the Pearson correlation between ***f***(*t*) and the measured FC. For each dataset, we can obtain the minimum, maximum, as well as the mean highest correlation values over all the subjects of the dataset. Then we compare the results using Laplacians with different order, including Laplaican of SC (*m*=0), Laplacian of *h*SC (*m*=1), as well as the Laplacian of the high-order of *h*SC (*h*^*m*^SC, *m*>1) to examine the performance changes with the number of orders.

For each dataset, the minimum, maximum, as well as the mean highest Pearson correlation values over all subjects using different Laplacians are summarized in Table S1. Results show that the mean highest Pearson correlations over the subjects for all the four datasets are increased by a big margin with the number of orders increasing when using Laplacian of *h*^*m*^SC and remain steady when the number of orders reaches 3 to 5 (*R*_90_= 0.8501 (*m*=3), *R*_246_= 0.8362 (*m*=3), *R*_998_=0.7769 (*m*=4), and *R*_2514_=0.7182 (*m*=5)), significantly exceeding the results of using the SC Laplacian (*R*_90_= 0.4112, *R*_246_= 0.3785, *R*_998_=0.3089, and *R*_2514_=0.2154; *m*=0) (Fig. 2), where all the *P* values of the Pearson correlations obtained in this paper satisfy *P* << 1*e* – 6. Fig. 3 shows the Pearson correlations using the Laplacian of *h*^*m*^SC with the largest order *m* for each subject of the four datasets varying with the parameter *βt*, respectively. It can be observed that the maximum Pearson correlation ranges from 0.7760 to 0.8861 for the 90-ROI dataset, 0.8012 to 0.8791 for the 246-ROI dataset, 0.7587 to 0.7972 for the 998-ROI dataset, and 0.6814 to 0.7575 for the 2514-ROI dataset, and the mean maximum correlation values for the four datasets take values 0.8501, 0.8362, 0.7769, and 0.7182, respectively. Fig. S2 demonstrates the spreading of the colour maps of FC along *h*^*m*^SC for the 90-ROI dataset using Brainnet viewer [36].

**Fig.2.**
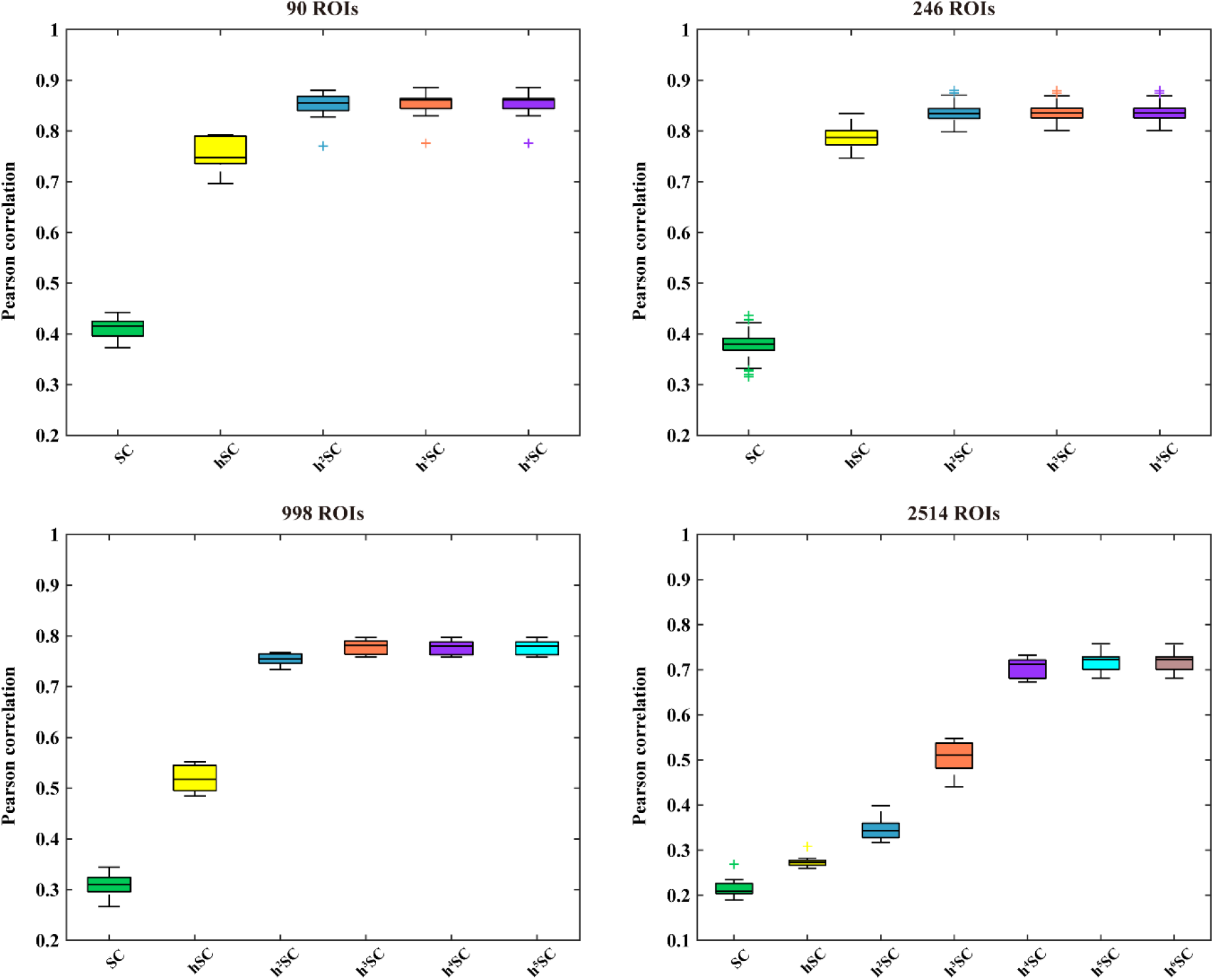
The maximum Pearson correlations distribution using Laplacian of *h*^*m*^SC (*m* = 0, referring to SC) for each subject of the four connectome datasets.

**Fig.3.**
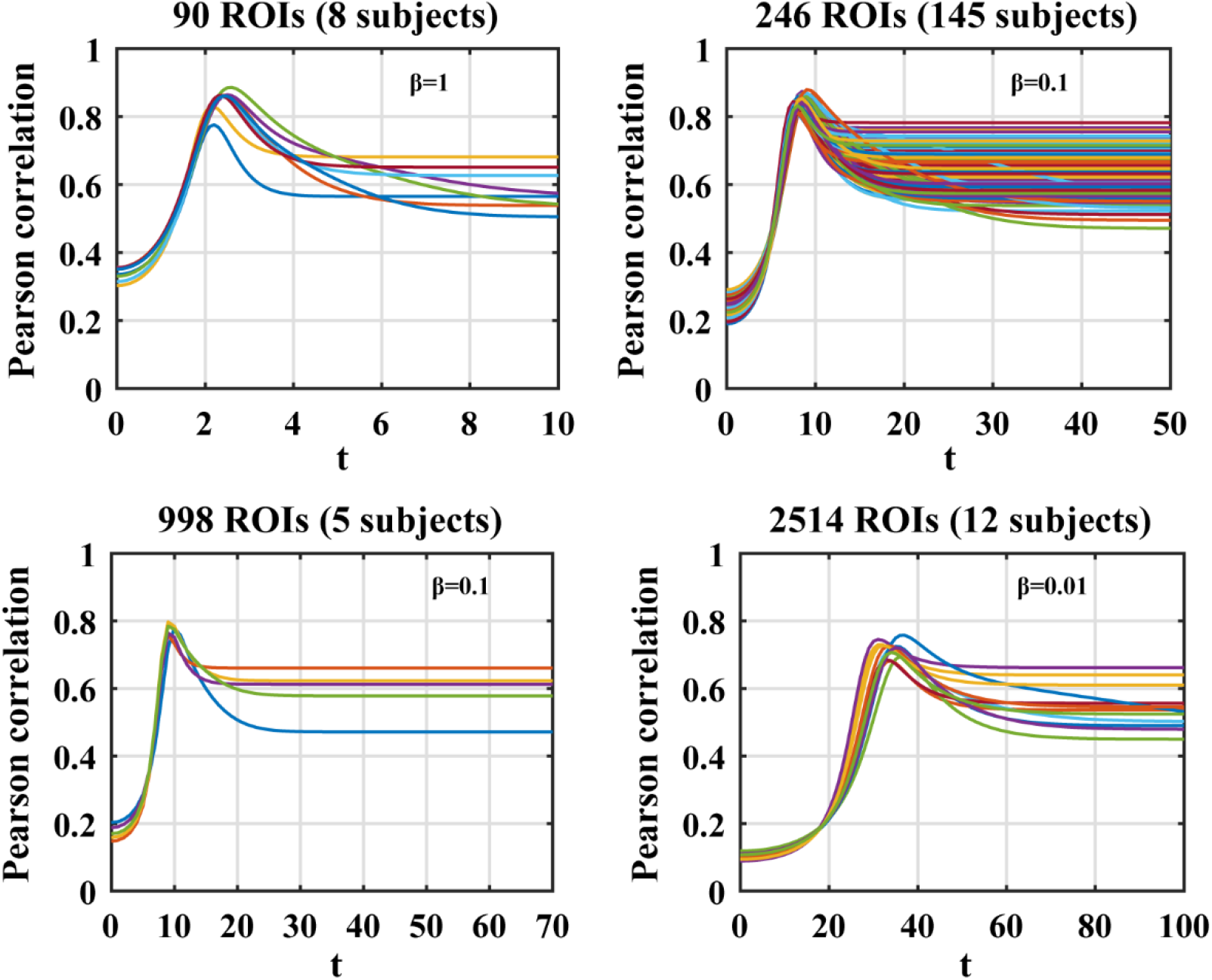
Pearson correlations using Laplacian of *h*^*m*^SC with the largest order *m* for each subject of the four datasets varying with the parameter *βt*

### 4.4 Structure-function mapping using the combined Laplacian of *h*^*m*^SC with SC

As described above, the brain cortical functions are constrained by the Laplacian of *h*SC varying linearly with time, leading to the varying of its eigenvectors representing harmonics and eigenvalues representing frequencies. Since the constantly varying *h*SC is rooted from the relatively fixed brain SC and, accordingly, we also apply the wave equation to characterize FC with the reconfigured Laplacian by combining the eigenvectors of *h*^*m*^SC Laplacian with the eigenvalues of SC Laplacian (denoted by *h*^*m*^SC+SC). For each dataset, the minimum, maximum, as well as the mean highest correlation values for all subjects using the combined Laplacian of *h*^*m*^SC and SC are summarized in Table S2. It can be clearly found that very high mean Pearson correlations are obtained across all the four datasets when binding the Laplacian eigenvectors of *h*^*m*^SC and the Laplacian eigenvalues of SC together after all the brain region are hyper-connected (*R*_90_= 0.8972 (*m*=3), *R*_246_= 0.9281 (*m*=3), *R*_998_=0.9097 (*m*=4), and *R*_2514_=0.9346 (*m*=5)) (Table S2, Fig.4), approaching the ceiling on the performance of SC-FC mappings with the models using eigen-decomposition [20]. Fig. 5 illustrates the Pearson correlations using the Laplacian of *h*^*m*^SC+SC with the largest order *m* for each subject of the four datasets varying with the parameter *βt*, respectively. It can be observed that the maximum Pearson correlation ranges from 0.8691 to 0.9146 for the 90-ROI dataset, 0.9120 to 0.9411 for the 246-ROI dataset, 0.9010 to 0.9146 for the 998-ROI dataset, and 0.9162 to 0.9487 for the 2514-ROI dataset, and the mean maximum correlation values for the four datasets are 0.8972, 0.9281, 0.9097, and 0.9346, respectively, far outperforming the cases using the Laplacian of *h*^*m*^SC alone.

**Fig.4.**
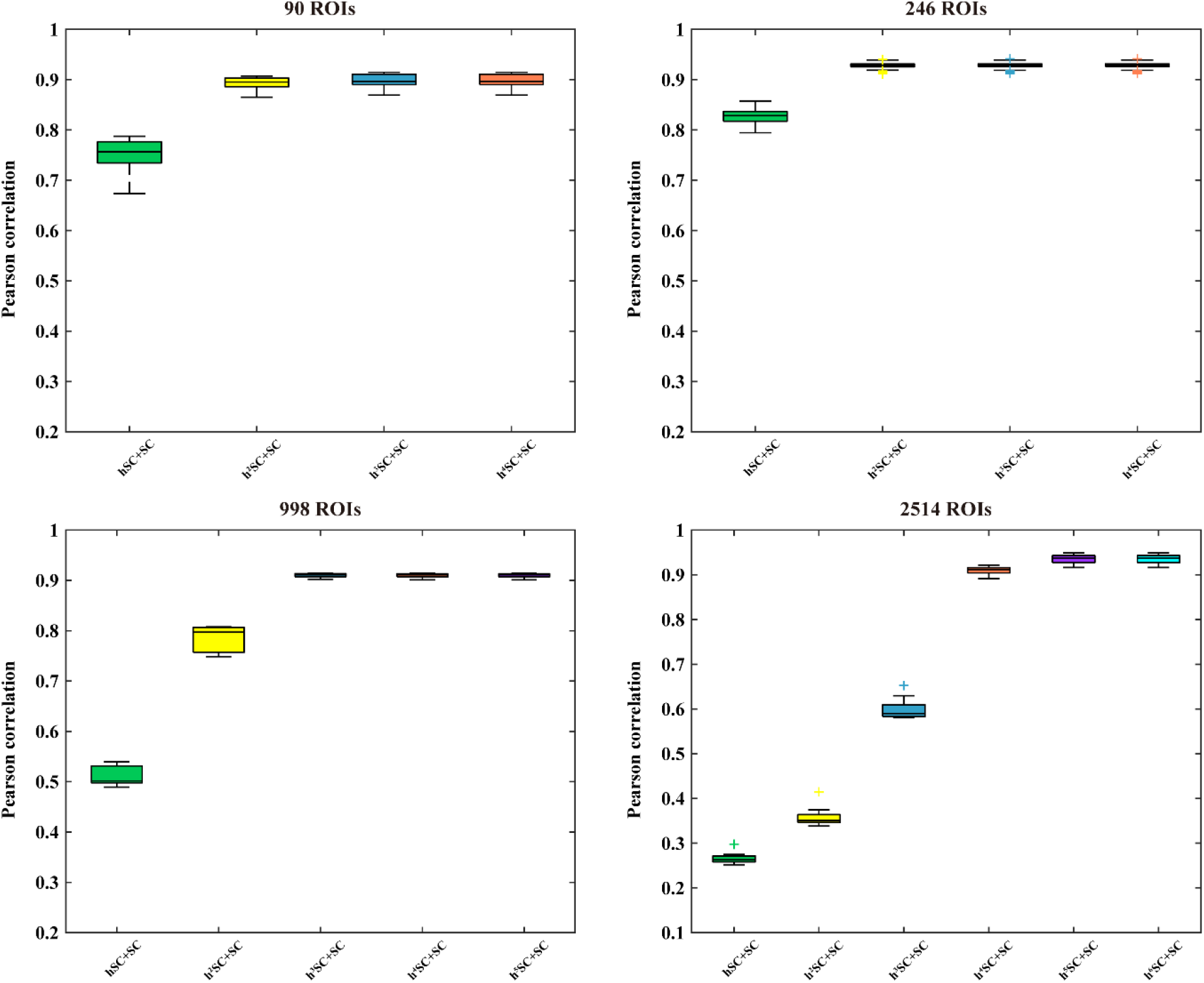
The maximum Pearson correlations distribution using the Laplacian of *h*^*m*^SC+SC (*m* > 0) for each subject of the four connectome datasets.

**Fig.5.**
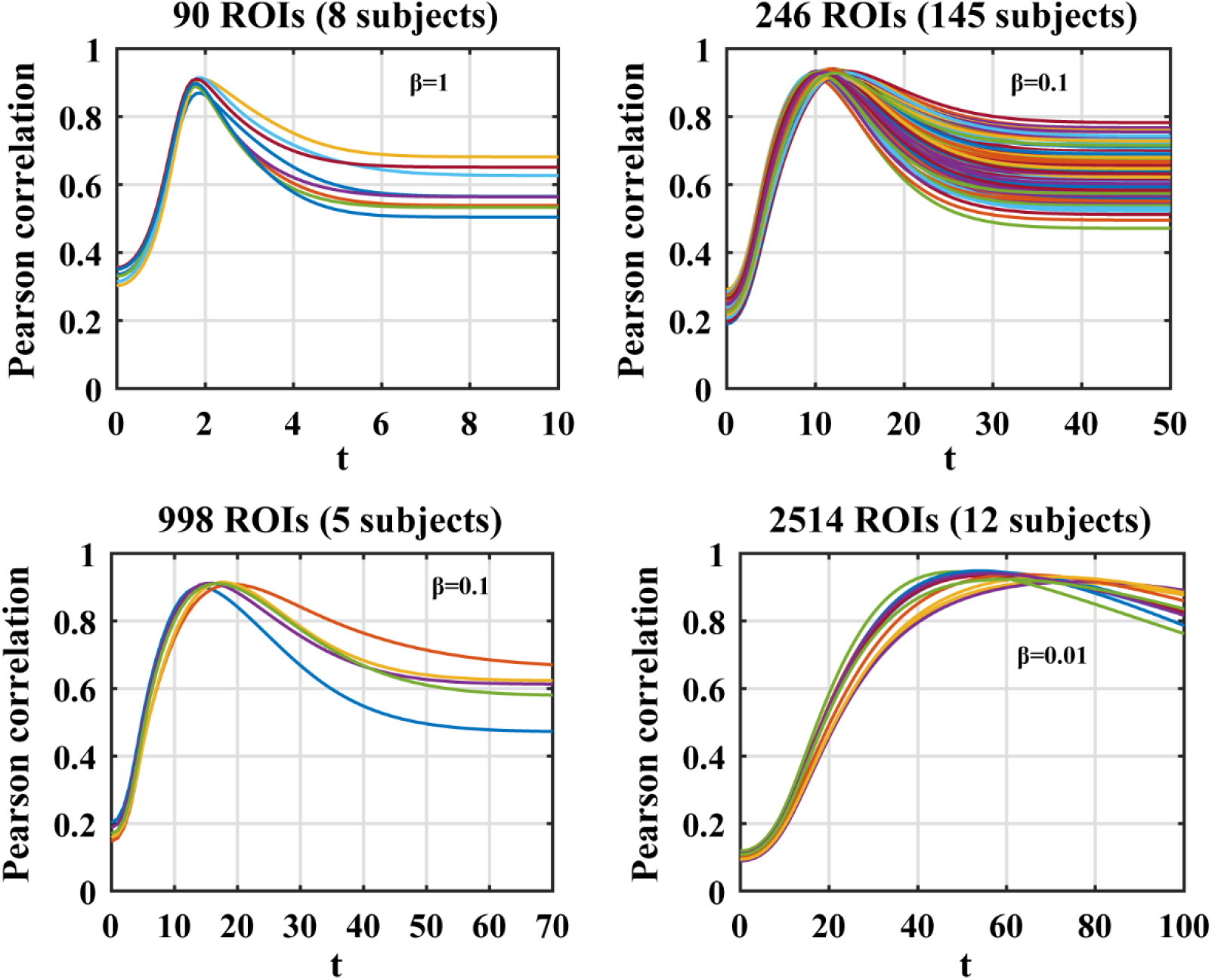
Pearson correlations using the Laplacian of *h*^*m*^SC+SC with the largest order *m* for each subject of the four datasets varying with the parameter *βt*

To demonstrate that the predicted FC (arising from *f*(*t*) at the time when the Pearson correlation value reaches the maximum) is not obtained by chance, for each dataset, we randomly permute the mean SC with 100 times, then for each scrambled SC, we extract the predicted FC and evaluate the performance through computing the histogram of the Pearson correlations between the mean predicted FC and the mean measured FC. For clear comparison, we also show the Pearson correlations between the mean measured FC and the mean predicted FC without permuting for the four datasets respectively (Fig. 6).

**Fig. 6.**
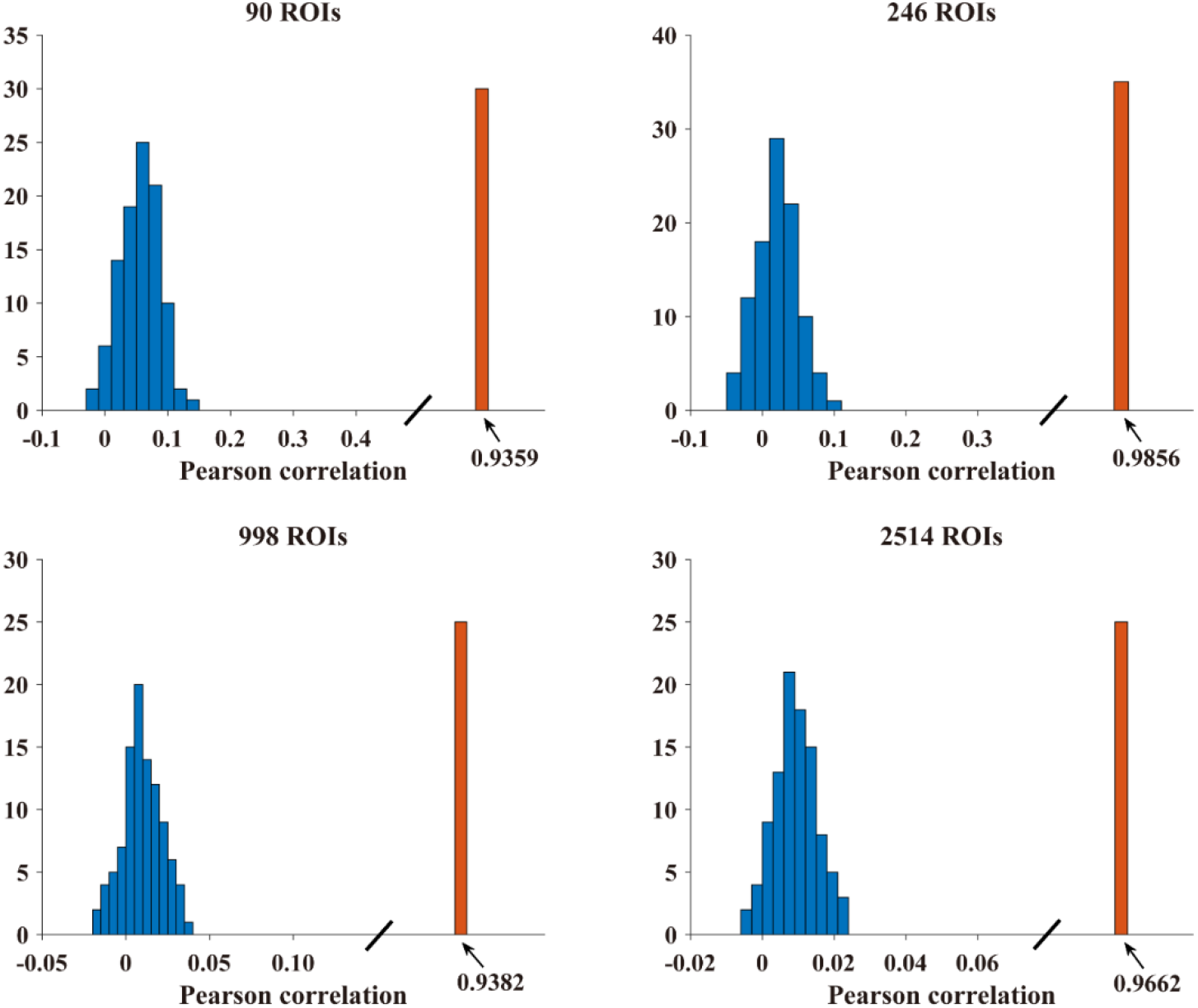
The histogram of the Pearson correlations between the mean measured FC and the mean predicted FC (blue) after permuting the mean SC with 100 times as well as the mean predicted FC without permuting (orange) for the four datasets.

## 5. Discussions

As well known, the Laplace operator plays a core role in elucidating the nature of heat, light, sound, electricity, magnetism, gravitation, etc. [37]. In this study, we extended the Laplace operator to the connectome Laplacian of human brain and established a link between the two, upon which we derived a wave equation to describe the relationship between human brain structure and function. We tested the proposed wave equation on four extensively studied experimental connectome datasets with increasing resolutions and the results show that the proposed wave equation is able to accurately and fast capture the relationship between brain SC and FC during rest.

Our major findings suggest that the dynamic couplings between brain regions at rest are largely regulated by the hypergraph Laplacian of SC varying in both time and space. The high-order hypergraph representation of SC may provide new insight into understanding the signal propagating in brain and help to shed light on the way of signals transmission between two distant brain regions without directly anatomical links. Previous studies have confirmed that the number of short connection patterns of length-2 paths is related to the presence of FC between structurally unconnected brain regions [2, 7–9, 38], including patterns of common efferents, common afferents, two-step serial relay, as well as the local detours along the shortest paths. Here we show that the high-order hypergraph representation of SC allows rapid signal transmission from one brain area to those anatomically unconnected brain areas via a third commonly connected area if these brain areas are contained in the same hyperedge (Fig. 1). As listed in Table S1, normally, only3-5 orders of mapping are required for the hypergraph to be fully connected for the four datasets including the one with 2514 ROIs, in which the SC of each subject is extremely sparse [35], being far less than the number of steps along the shortest paths [8].

Our results also suggest that the dynamic brain functions may favor the time-varying Laplacian with the eigenvectors of the Laplacian of *h*^*m*^SC and the eigenvalues of the Laplacian of SC (Fig. 5, Table S2). To put it another way, the resting brain cortical activities are more likely to propagate with harmonics (eigenvectors) emerging from *h*^*m*^SC while being constrained by the inherent frequencies (eigenvalues) of SC. These results are different from previously reported findings [20–22] in which the graph Laplacian of SC was regarded as time-invariant. It can be seen from Eq. (12) that, in our model, the hypergraph Laplacian of SC is linearly varied with time, thereby causing the cortical harmonics with higher frequencies (eigenvalues) attenuate rapidly during propagating until reach a steady state (Fig. 5). These results also imply that the human brain network works with critical frequencies when FC appears, after which the harmonic waves begin to taper off. This extends and supports the previously reported findings that the brain network operates at the edge of instability when FC emerges[39, 40]. Fig.7 demonstrates the fluctuation of the Pearson correlations between the ***f***(*t*) and the measured FC (Fig. 7 A) together with the trace change of each ***f***(*t*) (Fig. 7 B) over time. The trace changes are nearly the same for all subjects from the same dataset, with the initial value (*t*=0) being the number of brain regions and dropping down with time. When the correlation reaches a peak, the corresponding trace of ***f***(*t*), being the same as the sum of its eigenvalues (relating to certain harmonic frequencies) will fall below a critical value *λ*_*s*_(*λ*_*s*_ = 5 for datasets 90, 246, and 998 ROIs, and *λ*_*s*_ = 10 for 2514 ROIs) (Fig. 7 B). These critical values can be estimated by searching for the maximal curvature on the trace curve. Then we can determine the critical time *t*_*s*_ on ***f***(*t*) based on these critical trace values, which favors the prediction of FC using Eq. (15) directly. The mean Pearson correlations between the directly predicted FC and measured FC when using Laplacians of *h*^*m*^SC+SC are *R*_90_= 0.8921(*m*=3), *R*_246_= 0.9240 (*m*=3), *R*_998_=0.9054 (*m*=4), and *R*_2514_=0.9281(*m*=5), respectively, which are computed by averaging the correlations emerging at the same time for all subjects (i.e., *t*_*s*_ in Fig. 7) and a little lower than the highest mean correlations obtained by averaging the highest correlations arising at different time attributed to the inter-subject variability (Fig. 5, Table S2), thereby causing small deviations from the highest mean correlations. However, it should be noted that the predicted full FC patterns at the critical frequencies closely resemble the corresponding measured FC for all the four datasets (Fig. 8).

**Fig. 7.**
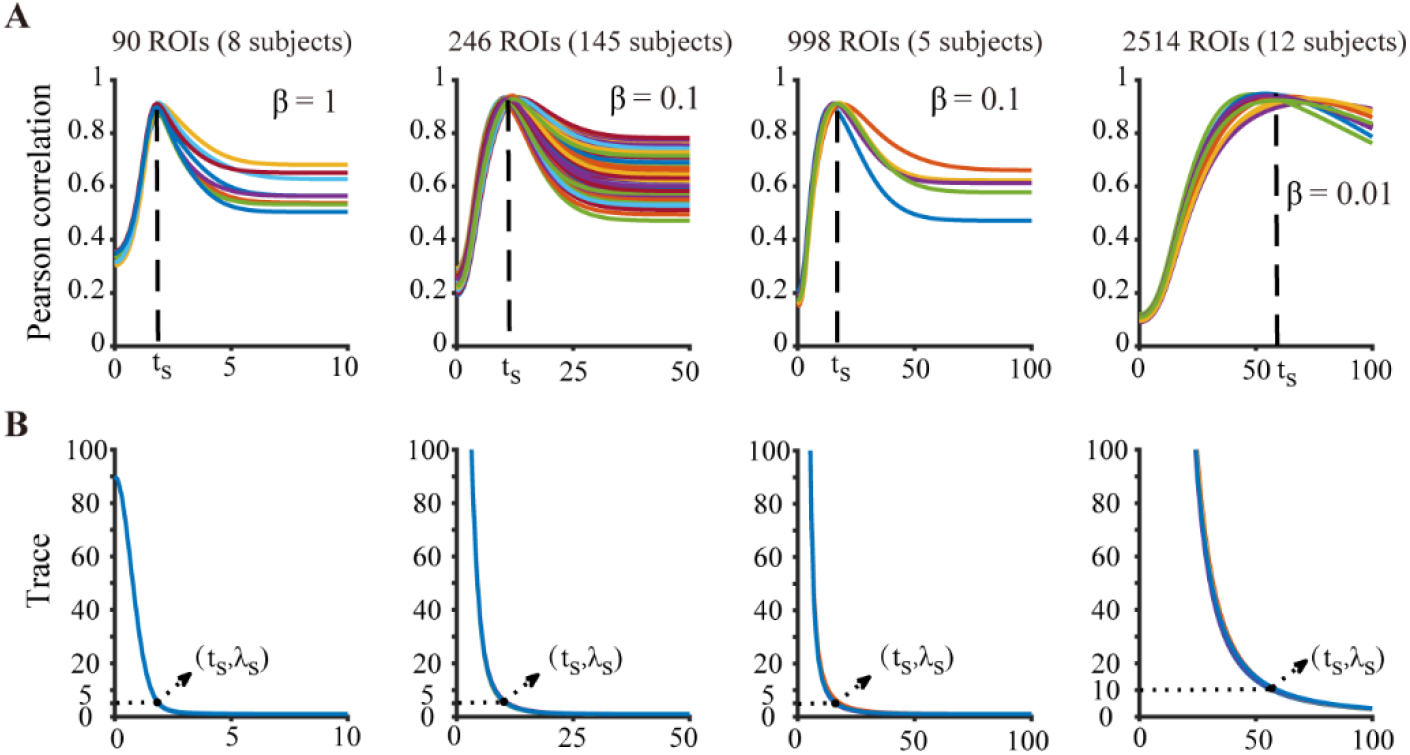
Illustration of the critical frequencies when FC appears for all the four datasets. **(A)** Pearson correlations between the instantaneous function (***f***(*t*)) and the measured FC over all subjects of the four datasets. **(B)** The corresponding trace changes of ***f***(*t*). The highest correlations (corresponding to the stationary FC) for each subject arise around the moment (*t*_*s*_) when the traces of ***f***(*t*) (corresponding to frequencies) fall below a critical value (*λ*_*s*_ = 5 for datasets 90, 246, and 998 ROIs, and *λ*_*s*_ = 10 for 2514 ROIs).

**Fig. 8.**
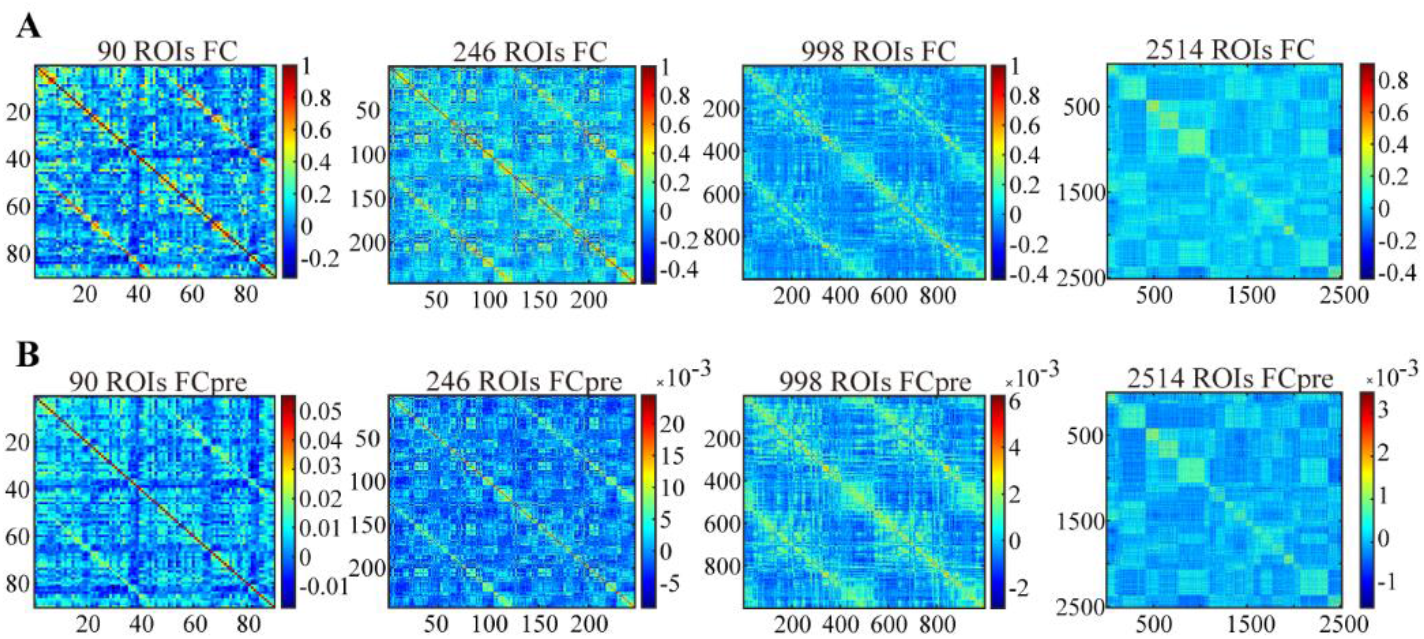
Comparison between the predicted FC matrices at the critical frequencies and the corresponding measured FC matrices for all datasets. **(A)** Mean measured FC matrices of the four datasets. **(B)** The corresponding mean predicted FC (FCpre). Mean Pearson correlations between the two are *R*_90_= 0.8921, *R*_246_= 0.9240, *R*_998_=0.9054, and *R*_2514_=0.9281 for the four datasets, respectively.

Another critical finding of our study is that the established brain wave equation is capable of modeling the anti-correlations between brain regions, which are generally neglected in previously reported models [6–9, 21]. These models usually exclude all the connections whose strength is smaller than a threshold including all the negative connections (functional anti-correlations) in FC. In our model, we argue that the negative correlations result from the long term depression (LDP) [32] occurring during signal transmission between brain regions. The lasting decrease of the connection strength between two brain regions may lead to the emergence of negative correlation between them. We observe that the predicted FC using the wave equation shows nearly the same signs as the measured FC by embedding the sign matrix extracted from the measured FC into the wave equation directly (Fig. S3). Very high mean Pearson correlations are obtained between the simulated negative FC and the true negative FC using Laplacians of *h*^*m*^SC+SC for all the four datasets (*R*_90_= 0.8606 (*m*=3), *R*_246_= 0.9825 (*m*=3), *R*_998_=0.9044 (*m*=4), and *R*_2514_=0.9378(*m*=5)).

However, it should be noted that, although the proposed wave equation is capable of modeling the negative correlations between brain regions, in the present study, we obtain the sign matrix from the measured FC directly due to the inability of dMRI to measure directed interactions between brain regions. Alternative approaches such as using graph neural fields [41] and deep learning modeling [42] or developing optimal noninvasive *in vivo* imaging techniques with high spatial and temporal resolution [43].

Taken together, it is worth highlighting that the established wave equation may bring to light how neural activities are coupled and propagate in brain. Although the current model lacks strong biological evidence, we have demonstrated its powerful performance on brain structure-function mapping using four extensively studied experimental connectome datasets and obtained compelling and exceptional results.

## Acknowledgments

Y. W. appreciates Richard M. Shiffrin and Olaf Sporns for their invaluable discussions on the research during the visit to Indiana University, Bloomington. The authors also cordially thank Farras Abdelnour for sharing the 90 ROIs connectome dataset together with the implementation codes of their network diffusion model, Olaf Sporns for sharing the 998 ROIs data, and Yong He for sharing the structural and diffusion MRI together with the resting-state fMRI data of 147 subjects from Beijing Normal University. Y. W. was supported by the China Scholarship Council (201306455001), the National Natural Science Foundation of P.R. China (Grant No.62072468). J. M. was supported by the Graduate Innovation Project of China University of Petroleum (East China) (Grant No. YCX2020095). X. C. was supported by the China Scholarship Council (201706450045) and the Fundamental Research Funds for the Central Universities (Grant No.16CX06050A).

## Author Contributions

Y. W. conceived and designed the research, wrote the manuscript. J. M., X. C. and C. D. performed the research.

## Compliance with ethics guidelines

The authors declare that they have no conflicts of interest or financial conflicts to disclose.

## Supplementary Materials for

**Fig. S1.**
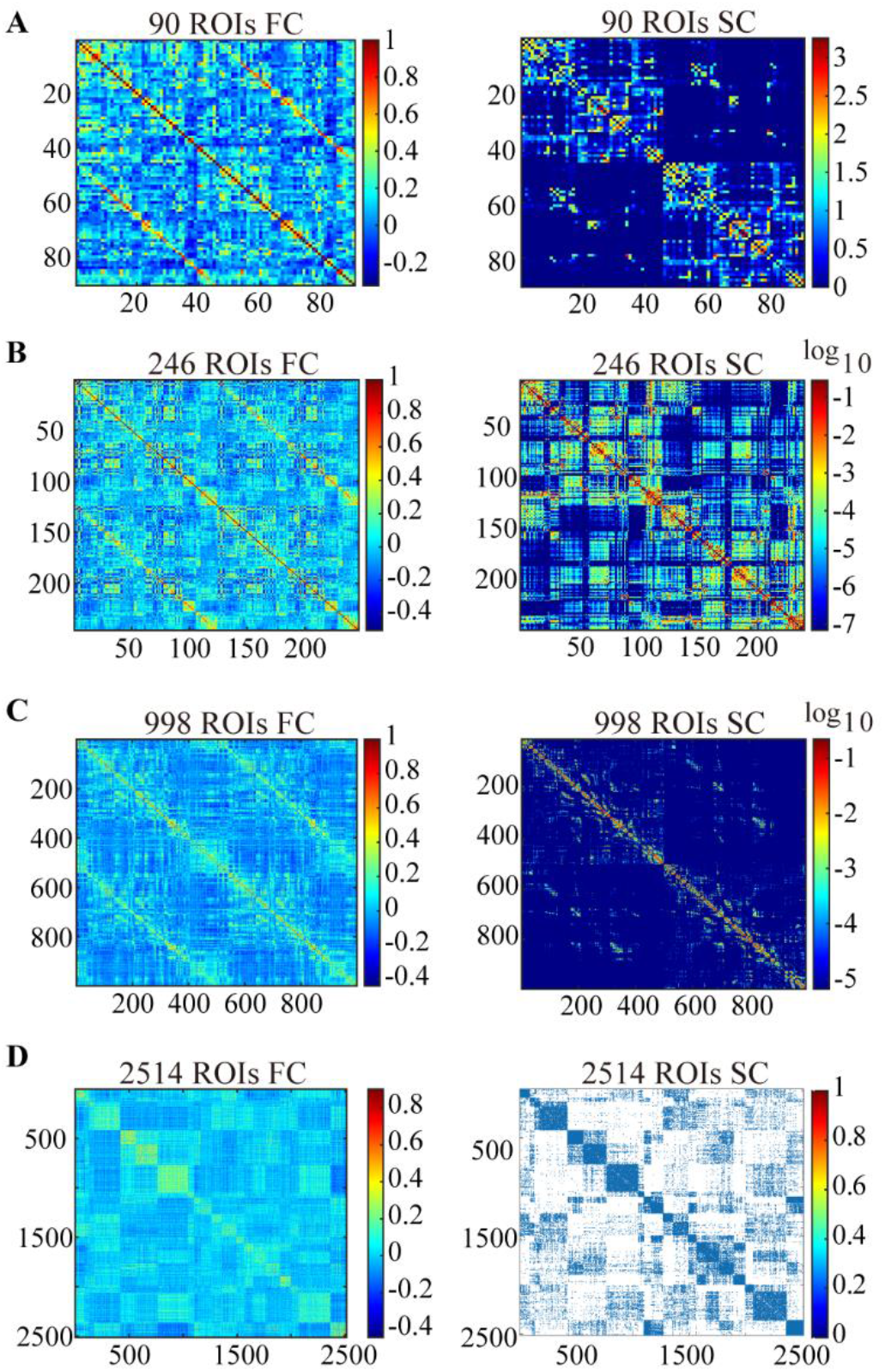
Mean measured FC and SC matrices of the four datasets. **(A)** 90 ROIs dataset, averaged across 8 individual participants. **(B)** 246 ROIs dataset, averaged across 145 individual participants. **(C)** 998 ROIs dataset, averaged across 5 individual participants. **(D)** 2514 ROIs dataset, averaged across 12 individual participants. The matrices’ elements of all the datasets except the 2514 ROIs are arranged such that the right hemisphere is in the upper left quadrant, left hemisphere in the lower right quadrant, and interhemispheric connections in the upper right and lower left quadrants. For clear visualization, the FC matrix of 2514 ROIs is modularized with hierarchical agglomerative clustering (HRC) and the SC matrix is binarized in the same order as those in the rsFC [35].

**Fig. S2.**
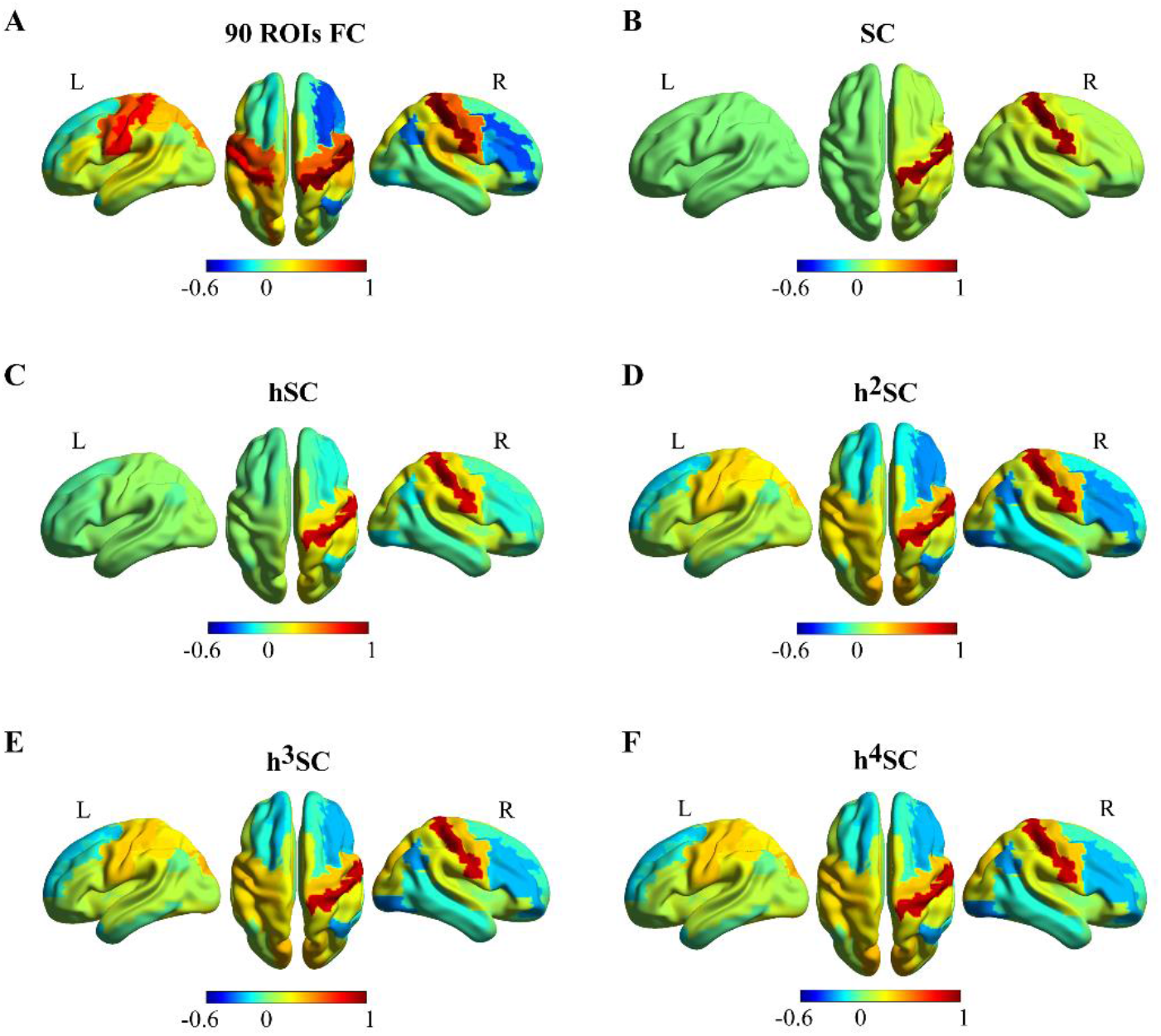
The spreading of the colour maps of FC along *h*^*m*^SC for the 90-ROI dataset. The color maps are obtained with seeding at right postcentral gyrus (PoCG.R) in the brain network. **(A)** the measured functional connectivity (FC) between the seed PoCG.R and the other regions. **(B)** the predicted FC between the seed PoCG.R and the other regions derived from the measured structural connectivity (SC). **(C), (D), (E)** and **(F)** show the predicted FCs between the seed PoCG.R and the other regions by the high-order of *h*SC (*h*^*m*^SC), with the number of order *m* = 1, 2, 3, and 4, respectively. Cold/warm colour indicates negative/positive functional connections.

**Fig. S3.**
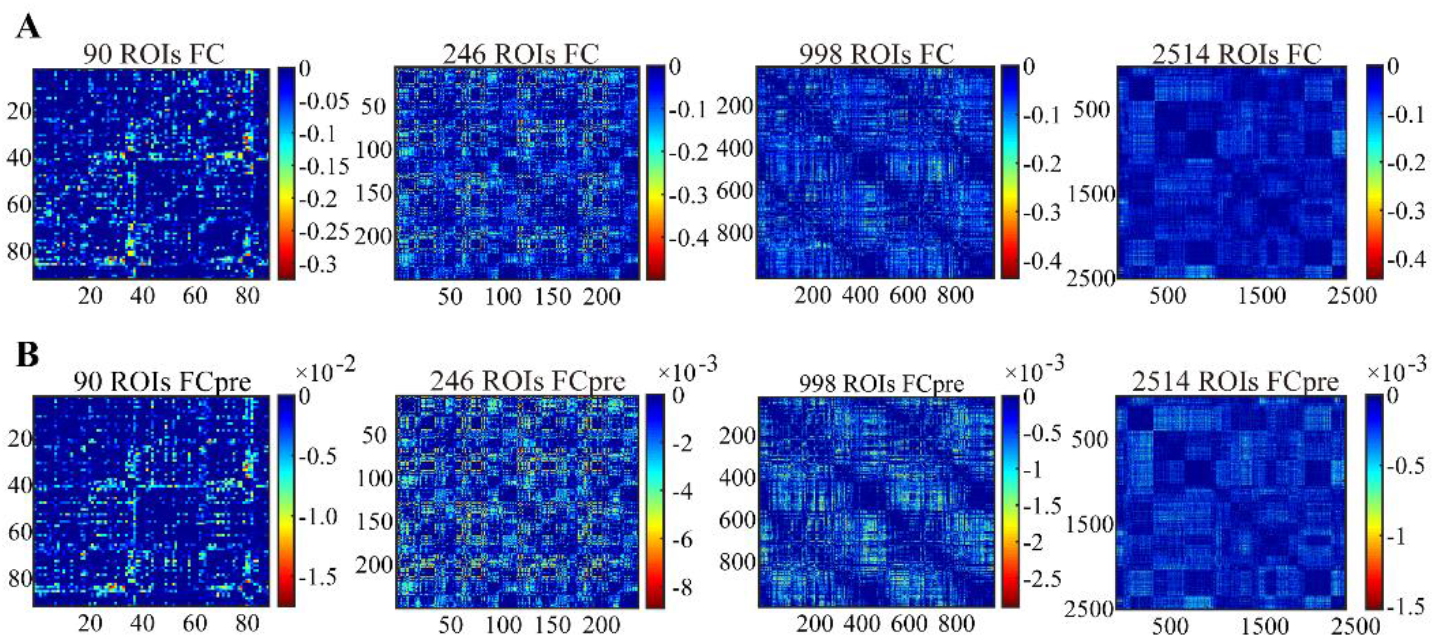
Comparison between the predicted negative FC matrices and the corresponding measured negative FC matrices for the four datasets. **(A)** Mean measured negative FC matrices of the four datasets. **(B)** The corresponding mean predicted negative FC (FCpre). Mean Pearson correlations are *R*_90_= 0.8606, *R*_246_= 0.9825, *R*_998_=0.9044, and *R*_2514_=0.9378, respectively.

**Table S1.**
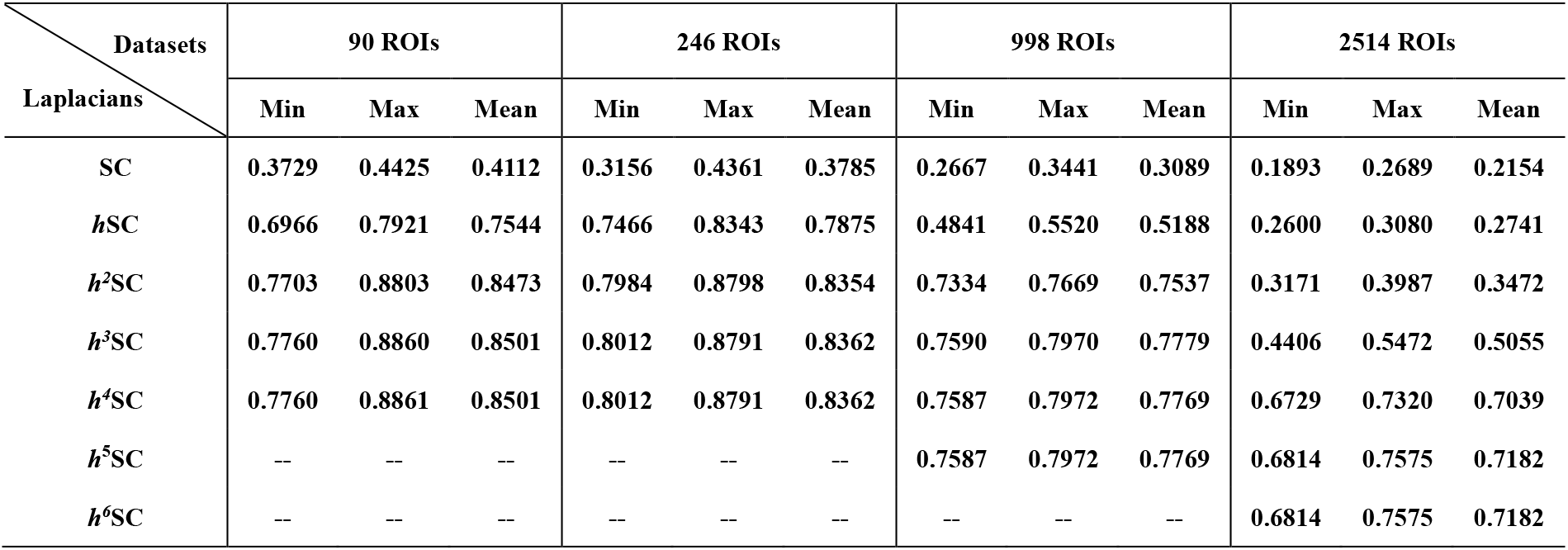
The maximum Pearson correlations distribution for the four datasets using Laplacians of *h*^*m*^SC.

| Datasets<br>Laplacians | 90 ROIs |  |  | 246 ROIs |  |  | 998 ROIs |  |  | 2514 ROIs |  |  |
| --- | --- | --- | --- | --- | --- | --- | --- | --- | --- | --- | --- | --- |
|  | Min | Max | Mean | Min | Max | Mean | Min | Max | Mean | Min | Max | Mean |
| SC | 0.3729 | 0.4425 | 0.4112 | 0.3156 | 0.4361 | 0.3785 | 0.2667 | 0.3441 | 0.3089 | 0.1893 | 0.2689 | 0.2154 |
| $h$SC | 0.6966 | 0.7921 | 0.7544 | 0.7466 | 0.8343 | 0.7875 | 0.4841 | 0.5520 | 0.5188 | 0.2600 | 0.3080 | 0.2741 |
| $h^2$SC | 0.7703 | 0.8803 | 0.8473 | 0.7984 | 0.8798 | 0.8354 | 0.7334 | 0.7669 | 0.7537 | 0.3171 | 0.3987 | 0.3472 |
| $h^3$SC | 0.7760 | 0.8860 | 0.8501 | 0.8012 | 0.8791 | 0.8362 | 0.7590 | 0.7970 | 0.7779 | 0.4406 | 0.5472 | 0.5055 |
| $h^4$SC | 0.7760 | 0.8861 | 0.8501 | 0.8012 | 0.8791 | 0.8362 | 0.7587 | 0.7972 | 0.7769 | 0.6729 | 0.7320 | 0.7039 |
| $h^5$SC | -- | -- | -- | -- | -- | -- | 0.7587 | 0.7972 | 0.7769 | 0.6814 | 0.7575 | 0.7182 |
| $h^6$SC | -- | -- | -- | -- | -- | -- | -- | -- | -- | 0.6814 | 0.7575 | 0.7182 |

**Table S2.** The maximum Pearson correlations distribution for the four datasets using Laplacians of *h*^*m*^SC+SC.

| Datasets<br>Laplacians | 90 ROIs |  |  | 246 ROIs |  |  | 998 ROIs |  |  | 2514 ROIs |  |  |
| --- | --- | --- | --- | --- | --- | --- | --- | --- | --- | --- | --- | --- |
|  | Min | Max | Mean | Min | Max | Mean | Min | Max | Mean | Min | Max | Mean |
| $h$SC+SC | 0.6735 | 0.7872 | 0.7494 | 0.7944 | 0.8573 | 0.8274 | 0.4889 | 0.5394 | 0.5117 | 0.2512 | 0.2974 | 0.2656 |
| $h^2$SC+SC | 0.8652 | 0.9070 | 0.8926 | 0.9116 | 0.9399 | 0.9281 | 0.7486 | 0.8080 | 0.7840 | 0.3383 | 0.4146 | 0.3576 |
| $h^3$SC+SC | 0.8691 | 0.9146 | 0.8972 | 0.9120 | 0.9411 | 0.9281 | 0.9023 | 0.9146 | 0.9100 | 0.5803 | 0.6526 | 0.5997 |
| $h^4$SC+SC | 0.8691 | 0.9146 | 0.8972 | 0.9120 | 0.9411 | 0.9281 | 0.9010 | 0.9146 | 0.9097 | 0.8915 | 0.9213 | 0.9095 |
| $h^5$SC+SC | -- | -- | -- | -- | -- | -- | 0.9010 | 0.9146 | 0.9097 | 0.9162 | 0.9487 | 0.9346 |
| $h^6$SC+SC | -- | -- | -- | -- | -- | -- | -- | -- | -- | 0.9162 | 0.9487 | 0.9346 |

### Supplementary Note 1: The derivation of the solution to the wave equation

As described in the main text, the wave equation can be expressed as

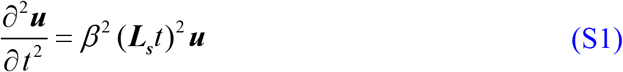

The solution is derived as follows.

Assume 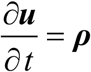, then 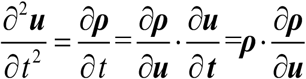, and then the wave equation can be rewritten as

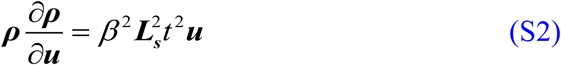

i.e., 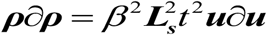, integrating on both sides gives,

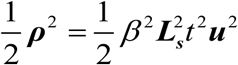

Considering the damping effect during the propagating of neural activity signals, the solution ***ρ*** can be expressed as

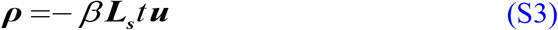

i.e., 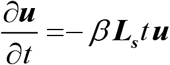, shifting ***u*** to the left side and ***t*** to the right side, we have

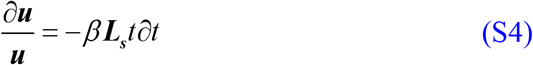

Integrating on both sides gives

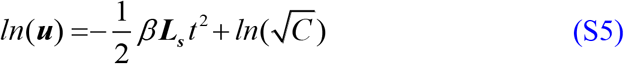

Then we obtain

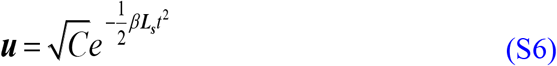

